# What are housekeeping genes?

**DOI:** 10.1101/2021.02.14.431149

**Authors:** Chintan J. Joshi, Wenfan Ke, Anna Drangowska-Way, Eyleen J. O’Rourke, Nathan E. Lewis

**Affiliations:** Department of Pediatrics, University of California, San Diego, School of Medicine, La Jolla, CA 92093; Department of Biology and Cell Biology, University of Virginia, Charlottesville, VA 22903; Novo Nordisk Foundation Center for Biosustainability at the University of California, San Diego, School of Medicine, La Jolla, CA 92093; Department of Bioengineering, University of California, San Diego, La Jolla, CA 92093; National Biologics Facility, Technical University of Denmark

**Author notes:** Corresponding authors: Name: Nathan E. Lewis, Name: Eyleen J. O’Rourke.

**Keywords:** omics data, systems biology, housekeeping genes

## Abstract

The concept of “housekeeping genes” has been used for four decades but remains loosely defined. Housekeeping genes are commonly described as “essential for cellular existence regardless of their specific function in the tissue or organism”, and “stably expressed irrespective of tissue type, developmental stage, cell cycle state, or external signal”. However, experimental support for the tenet that gene essentiality is linked to stable expression across cell types, conditions, and organisms has been limited. Here we use genome-scale functional genomic screens, bulk and single-cell sequencing technologies to test this link and optimize a quantitative and experimentally validated definition of housekeeping gene. Using the optimized definition, we identify, characterize, and provide as resources, housekeeping gene lists extracted from several human datasets, and 12 other animal species that include primates, chicken, and *C. elegans*. We find that stably expressed genes are not necessarily essential, and that the individual genes that are essential and stably expressed can considerably differ across organisms; yet the pathways enriched among these genes are conserved. Further, the level of conservation of housekeeping genes across the analyzed organisms captures their taxonomic groups, showing evolutionary relevance for our definition. Therefore, we here present a quantitative and experimentally validated definition of housekeeping genes that can contribute to better understanding of their unique biological and evolutionary characteristics.

## Introduction

The concept of housekeeping genes has aided theoretical and applied biology and the study of evolution. At the grand level, housekeeping genes can be defined as the minimal set of genes required to sustain life^1^. At the practical level, they can be defined as genes stably expressed in all cells of an organism under normal conditions irrespective of tissue type, developmental stage, cell cycle state, or external signal, or as markers of an organism’s healthy biological state^2^. At the evolutionary level, they may allow us to define species and higher taxa-specific genomic features^3–5^ and gene functions^5–7^ that may drive conservation or change. Thus, knowledge of housekeeping genes can significantly contribute to explorative, basic, and translational studies. However, despite the fundamental and translational utility of the concept of housekeeping gene, for over four decades their definition has remained axiomatic.

Housekeeping genes are often defined as being stably expressed in all cells and conditions^8^. However, expression stability is often tested in a handful of cell types and conditions, and then inferred to be stable across most cell types and conditions. Further, housekeeping genes have been proposed to be essential^2^, belong to cellular maintenance pathways^8–12^ and be conserved^5,6^. However, expression stability (similar expression across cell types and conditions), function (e.g., belonging to cellular maintenance), essentiality (loss-of-function is lethal), and conservation (in this context, stably expressed and essential across taxa) are four very different properties of a gene (Figure 1). No study has yet formally tested the relationships or potential linkage between these four properties. For example, it is unknown if or how the expression pattern of a gene informs its essentiality or ‘housekeepingness’ conservation. Thus, experimental support for the implied association between the properties defining housekeeping genes remains to be reported.

**Figure 1.**
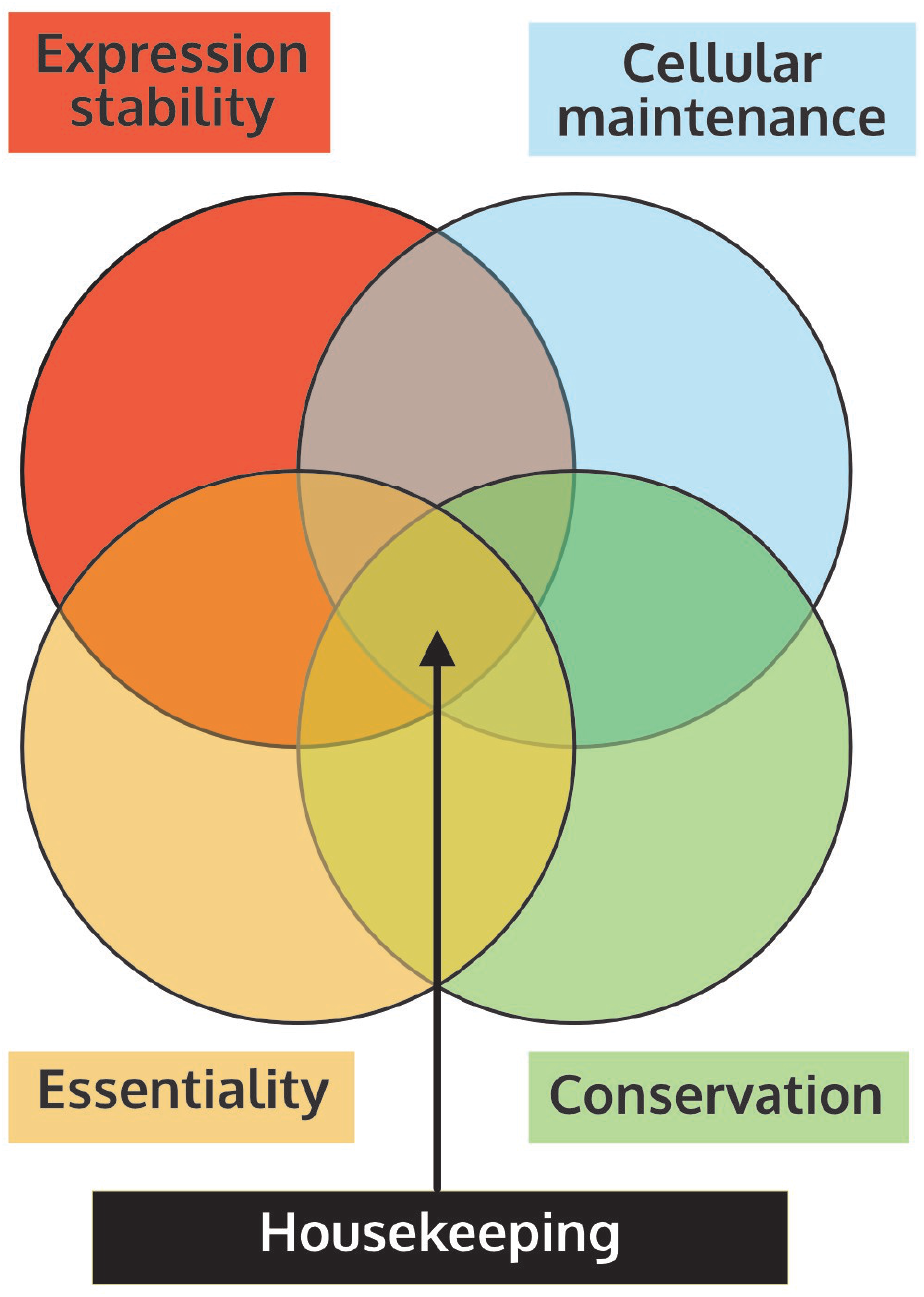
The relationship between the axiomatically assigned four properties of housekeeping genes remains unexplored.

Analysis of large-scale ‘omics’ data is now commonplace^13^, applied to a gamut of questions^14–17^ and organisms^18–25^. Further, the availability of single-cell transcriptomics and functional genomics studies for several species is rife with opportunities to study biological principles within and across organisms^23,26,27^, including developing a more objective and experimentally validated definition of housekeeping gene.

Predefining housekeeping genes for an organism would bring several potential benefits. At the experimental level, it can save in troubleshooting for the identification and validation of mRNA expression controls in difficult cell types and unique samples (e.g., seasonal wild organisms or patient biopsies) analyzed via transcriptomics^12^ and quantitative real-time PCR (qRT-PCR)^28^, and it can provide more robust ways to normalize the growing number of single-cell RNAseq studies. At the more fundamental level, it is important to define the minimum number of genes necessary to sustain life and if and how these genes or their characteristics change across taxa.

Recently, we presented StanDep^29^, a pipeline for constructing context-specific metabolic network models. StanDep effectively captures metabolic housekeeping genes, defined as genes expressed in most of the analyzed contexts (e.g., tissues, cell lines, etc). The capacity of StanDep to capture housekeeping genes – defined by expression stability – can be attributed to its effectiveness at capturing transcriptomic variability between different samples. Other recent efforts have been made to quantitatively define housekeeping genes^8,28,30,31^. A particularly powerful approach is a mathematical framework called GeneGini^7,30,31^, which leverages the Gini coefficient (G_C_) – a statistical metric quantifying inequality among groups^32^. G_C_ varies from 0 to 1. In economics, lower Gini coefficients mean lower income inequality. Similarly, in the framework of GeneGini, the G_C_ of a gene is proportional to the inequality in its expression across samples^30^. Therefore, genes with a low G_C_ (referred to as Gini genes here on) are stably expressed and could be considered housekeeping genes according to the property of stability. However, many questions remain about Gini genes. Which are the cellular functions carried out by Gini genes? Are Gini genes essential? Do Gini genes retain their housekeeping status across species? Answers to these questions are central to validate or reject the hypothesis that all four properties – expression stability, basic cellular function, essentiality, and conservation – are linked, and hence should be used to define housekeeping genes.

Here, we used the G_C_ approach to identify stably expressed genes (Gini genes) across human tissues and cell lines, and across cell types of lower organisms studied using single-cell transcriptomics^19,22,33–36^. We show that, indeed, G_C_ values are highly correlated across human datasets, supporting the existence of a subset of genes stably expressed across cell types and conditions. Then, we compare the properties of stability (G_C_ coefficient) to the property of essentiality, as defined by *in vitro* or *in vivo* functional genomics studies. Published CRISPR-Cas9 essentiality screens of Chinese-hamster ovary cells (CHO)^37^ and human cell lines^38–40^, and new whole-animal essentiality RNAi screening of *C. elegans* done as part of this study, show that essential genes tend to have lower G_C_, suggesting that stably expressed genes are more likely to be essential. Nevertheless, not all stably expressed genes are essential. Further, genes that are stably expressed and essential are associated with GO terms enriched in basic cellular functions, and are more likely to be conserved. Therefore, although the four properties informally used to define housekeeping genes are more likely than not to associate with each other, they are not strictly linked. Thus, our analysis provides an experimentally quantitative definition of housekeeping gene, and it establishes a foundation to start asking the most interesting questions including which is the minimum set of genes necessary to sustain life, and if and how these genes change across taxa.

## Results

### G_C_-defined housekeeping genes are stably expressed across samples of the same species

To quantify G_C_ across genes and test whether the G_C_ captures the first property of housekeeping genes (i.e., stable expression across cell types and conditions), we calculated the G_C_ for all genes in the Genotype-Tissue Expression (GTEx)^34^ and Human Protein Atlas (HPA)^22^ tissue datasets, and all genes in the NCI-60 Cancer datasets from CellMiner and Klijn et al. We used the 3688 housekeeping genes published by Eisenberg and Levanon^8^ as our benchmark gene set. A total of 15687 genes were present in HPA and GTEx, 16052 genes were present in both NCI-60 cancer datasets, and 14327 genes were present in all 4 datasets. From each dataset, we extracted the 3688 genes that had the lowest G_C_, this sample-independent gene set allows us to account for the different distribution shapes and number of genes in each dataset (Fig. S1A).

The observations follow: 1) the Gini genes obtained from combining HPA and GTEx (human tissues) covered 81.4% of the 3688 Eisenberg and Levanon housekeeping genes and 2) the NCI-60 cancer datasets covered 69.8% of the 3688 housekeeping genes (**Figure 2A**). Lower accuracy in the human cancer datasets relative to the human tissue datasets is expected as Eisenberg and Levanon housekeeping genes were defined using healthy human tissue transcriptomics^8^. Nevertheless, the G_C_ for genes present in any given pair of datasets were highly correlated, and more so for datasets of the same sample types (healthy or cancer samples)(**Figure 2B**). Further, we found that for all datasets, Eisenberg and Levanon’s housekeeping genes had low G_C_ values (Fig. S1B), and cross comparison of G_C_s revealed that the median G_C_ of these genes was very similar across samples (Fig. S2).

**Figure 2.**
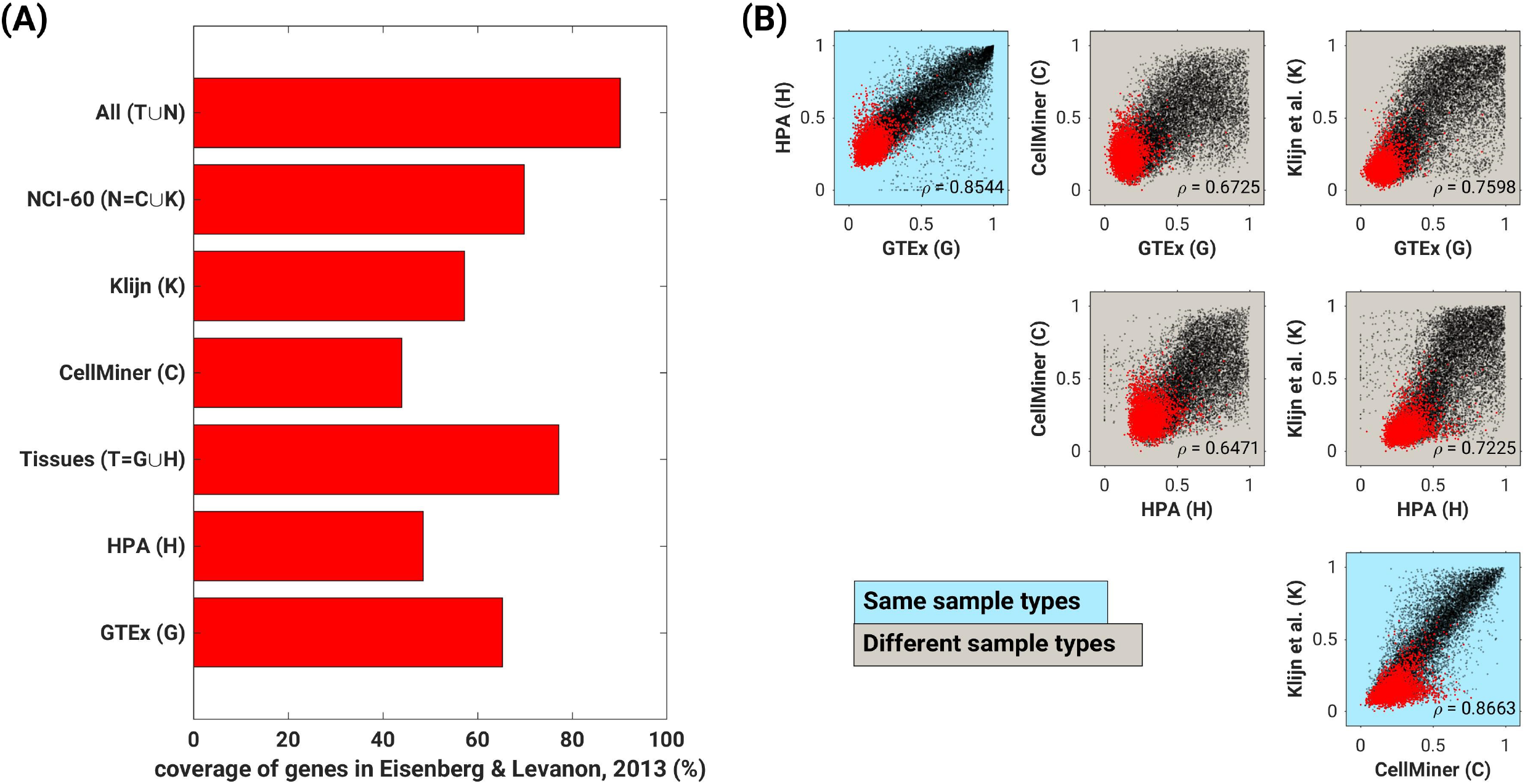
Gini coefficient captures the first property of housekeeping genes: stable expression. **(A)** Coverage of the 3688 Eisenberg and Levanon housekeeping genes^8^ among the 3688 genes with lowest Gini coefficients within each dataset. **(B)** Gini coefficients of individual genes are highly correlated across human datasets regardless of sample type; however, correlations within sample types are tighter. Dots represent unique genes. Red specifically depicts housekeeping genes identified by Eisenberg and Levanon^8^.

Although the Gc is informative regarding consistency of expression across samples, it does not inform about the actual level of expression of genes. To investigate whether housekeeping genes are represented across the range of expression or limited to one or other extreme, we used the GTEx^34^ and HPA datasets^22^ to calculate the median expression levels of the 3688 housekeeping genes previously identified by Eisenberg and Levanon^8^. We observed low median expression for the 3688 genes (**Figure 3 A & B**). Specifically, the median expression value within the 25^th^, 50^th^ and 75^th^ percentiles of the HPA dataset, were 12.2, 22.1, and 41.5 transcripts per million (TPM), respectively; whereas within the GTEx data set, the 25^th^, 50^th^ and 75^th^ percentile values were 14.2, 24.7, and 46.4 TPM, respectively.

**Figure 3.**
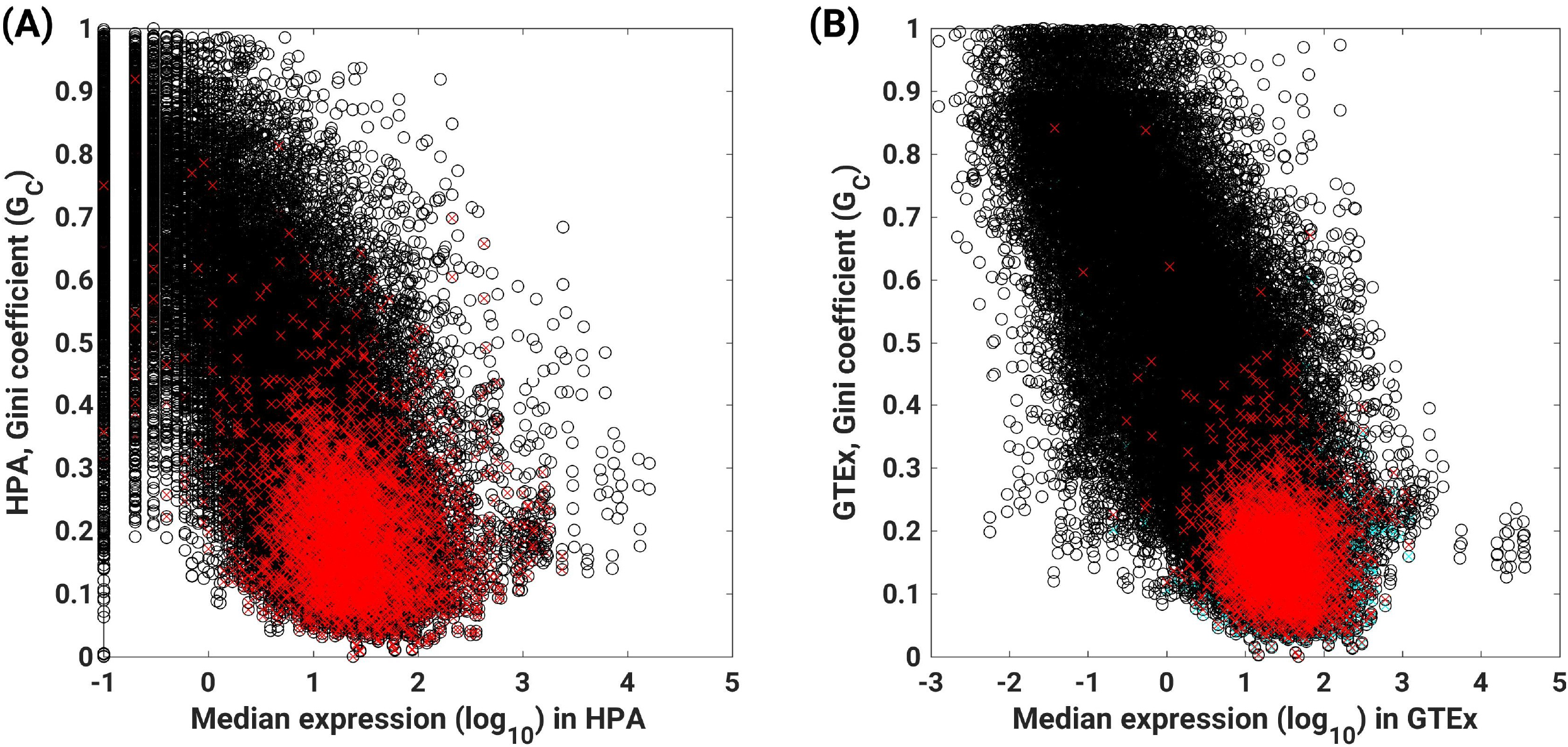
Housekeeping genes are expressed at relatively low levels. **(A-B)** Comparison of median expression levels and Gini coefficients calculated for the **(C)** HPA and (B) GTEx datasets. Dots represent unique genes. Red specifically depicts housekeeping genes identified by Eisenberg and Levanon^8^.

In summary, genes with low G_C_ in one sample type are the most likely to have low G_Cs_ in other sample types or datasets, suggesting that G_C_ captures the property of expression stability, at least within a given species. Additionally, distinct from current axiom, we found that the majority of the housekeeping genes are expressed at relatively low levels.

### G_C_-defined housekeeping genes are enriched in cellular maintenance functions

The second property commonly associated to the concept of housekeeping genes is that these genes are involved in basic cellular maintenance functions^8–12^. To test this axiom, we performed GO term enrichment analysis on the 3688 genes with lowest G_C_ identified for each dataset as described above.

First, to define the baseline GO terms, we extracted all GO terms enriched in the complete HPA, GTEx, CellMiner and Klijn *et al* samples. We found 1189 different GO terms that were enriched in at least one dataset (Fig. S3, Table S1). To minimize the confounding effect of using different numbers of subject or query genes for the hypergeometric test, we defined as background frequency of a GO term the representation of the compounded 1189 GO terms among all the genes of each dataset. We found that the representation of the 1189 GO terms was highly correlated across datasets (ρ_mean_ = 0.93 across 6 pairs of datasets). Then, we performed the enrichment analysis of the 3688 genes with lowest G_C_ identified in each dataset (tissue and cancer cell line datasets); details about hypergeometric test can be found in the Methods section.

We found 121 GO terms commonly enriched in all four datasets. The top GO terms with highest coverage were functions classically considered basic cellular maintenance including cell cycle, regulation of mRNA stability, protein folding, protein stabilization, and protein transport, among others. By contrast, the GO terms that were not enriched were related to stress or other context-specific responses including positive regulation of viral life cycle, response to mitochondrial depolarization, antifungal humoral response, and cellular response to misfolded protein, among others. The nature of the GO terms suggest that the GO terms most enriched among genes with low G_C_ relate to essential growth functions whereas the least enriched terms relate to regeneration and adaptation to stress rather than generation of new cells.

We also found sample type-specific GO terms; for instance, a GO term that was enriched in tissue data but not in cancer or vice-versa. We found 77 GO terms enriched exclusively in tissue datasets. The top 10 tissue-specific terms included positive regulation of mitochondrial translation and calcium ion transmembrane transport, branched-chain amino acid (BCAA) catabolic process, 7-methylguanosine mRNA capping, and NIK/NF-κB signaling. Excitingly, mitochondrial translation has been identified as a target for various cancers^41,42^, BCAA have been identified as an essential nutrient for cancer growth^43^. Further, enrichment of NF-κB highlights how a consistently expressed pathway controls expression of tissue specific genes. Thus, at least 3 out of 4 of the GO terms we find enriched in healthy tissues but not enriched in cancer cells correspond to biological functions known to be impaired or dysregulated in several cancers, suggesting that analyses of sample-specific genes with low Gc could contribute to identify disease-specific genes that could be become targets to treat or cure these diseases. Thus, our list reinforces the notion of enrichment of cellular maintenance functions.

We also found 70 GO terms enriched only in the cancer datasets, suggesting that cancer cells have housekeeping genes which are different than healthy tissues. These 70 cancer-only GO terms included mitotic cell cycle and replication fork machinery, GO terms associated with the generation of new cells. Further, the overall –as opposed to exclusive– coverage of GO terms enriched in cancer datasets was higher than the overall coverage of GO terms enriched in tissue datasets (Fig. S4), supporting the notion that the oncogenic state is an overall gain of function.

In summary, the data show that low G_C_ is associated with cell maintenance or cell generation activities, supporting the notion that housekeeping genes defined using the G_C_ would predominantly be involved in basic cellular functions. Beyond, the G_C_ identifies as housekeeping genes, core genes that are fundamental to homeostasis.

### Housekeeping genes are essential

A third property assigned to housekeeping genes is essentiality. However, this has only been validated for a handful of genes circumstantially found to be essential. Since no study has systematically tested whether essential genes are overrepresented among genes with stable expression, we here establish whether genes stably expressed (low Gc) are enriched for essential genes using functional-genomics studies in mammalian cells publicly available, as well as an *in vivo C. elegans* essentiality screen performed specifically for this study.

Essentiality in cancer cell line datasets was extracted from Depmap^38,39^ as a CRISPR guide-RNA score (log-fold change of guide-RNA). For essentiality scores, we used a published functional genomics screen that identified 338 essential genes in CHO cells^37^ (Table S5). The accession IDs of transcriptomic data for CHO cells are listed in Table S6. We found that the 2796 genes that were essential in all 20 cancer cell lines had lower G_C_ compared to all the genes combined, when calculated using transcriptomics data for the same 20 cell lines from Klijn et al. and CellMiner (Figure 4A and B). The same was true for 338 essential genes in CHO cells (Figure 4C). To facilitate comparison of GO terms between essential genes and genes with low G_C_ values, 2796 genes in cancer cells and 338 genes in CHO cells with lowest G_C_ values were chosen. This analysis of cancer cell lines revealed that coverage of GO terms for the essential genes is correlated with that of same number of low G_C_ genes identified using Klijn et al. (Figure 4D, yellow; ρ = 0.8907) and CellMiner (Figure 4D, blue; ρ = 0.8557). A similar comparison between CHO essential genes and genes with low G_C_ also resulted in a high correlation of ρ = 0.7055 between low G_C_ and essentiality (Figure 4D, green). Together these results suggest that Gini genes and essential genes show the same distribution, and hence, are likely largely overlapping.

**Figure 4.**
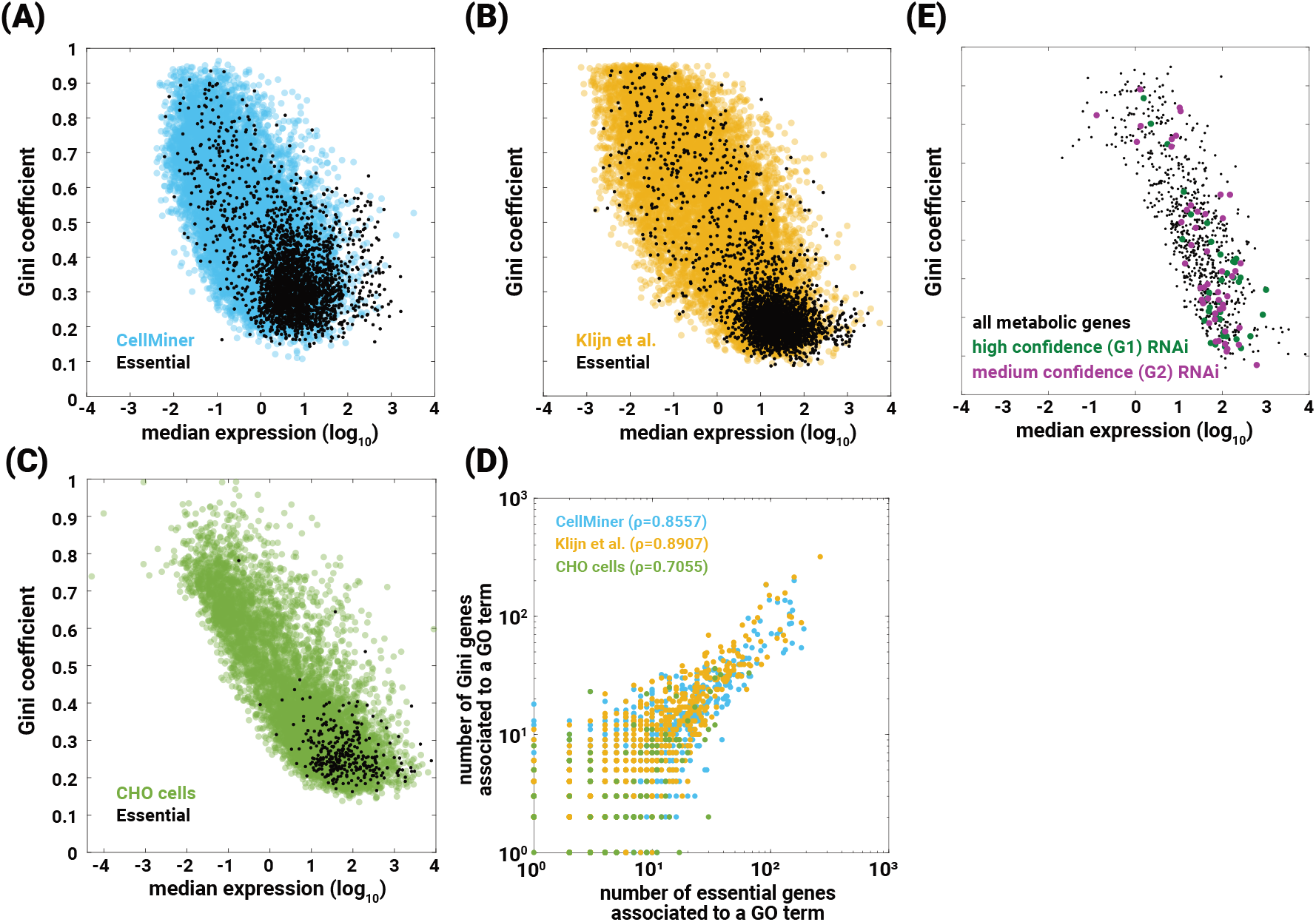
Gini genes are essential. Gini coefficients of essential genes (black dots) compared to the complete (A) CellMiner and (B) Klijn et al. cancer datasets, and (C) CHO datasets. The 2800 genes essential in the 20 cancer cell lines were extracted from DepMap^38,39^, and the 338 CHO essential genes were extracted from Xiong et al. (D) GO term coverage of essential genes and that of Gini genes from CellMiner (blue, 0.8557), Klijn et al. (yellow, 0.8907), and CHO (green, 0.7055) are correlated. The slightly lower correlation in CHO cells is likely due to the fewer number of essential genes reported in the CHO screen. (E) High- and medium-confidence essential genes in healthy *C. elegans* have significantly lower G_C_ than non-essential genes (p = 4.53 x 10^-5^).

Gene essentiality is: 1) health-status dependent, 2) context-dependent, and 3) subject to buffering from higher-level cell-cell interactions. Thus, since cell lines studied here are immortalized cultured cells devoid of their physiological and multicellular context, we investigated the correlation between G_C_ and gene essentiality in a healthy living animal model.

We defined the G_C_ for *C. elegans* genes using a whole-body single-cell transcriptomics dataset^18^. We obtained essentiality scores from our *in vivo* RNAi screen for all 1535 predicted metabolic genes in the worm (Table S4). A gene was considered essential in *C. elegans* if after rearing hatchlings on dsRNA-delivering *E. coli* for 5 days at 25°C (*E. coli* is the standard lab food source for *C. elegans*) animals were arrested at a pre-adulthood stage (control WT animals are gravid adults at this time). Three relevant phenotypic classes were observed in RNAi-treated *C. elegans:* 1) High-confidence essential, RNAi-treated animals arrested in ≥5 out of the 6 independent RNAi treatments against that gene; 2) Medium-confidence essential, RNAi-treated animals arrested in 3 or 4 out of the 6 independent RNAi treatments; and 3) Wild-type; see Supplementary Methods for further details. When testing the association between G_C_ and gene essentiality, we found that, similar to human and hamster diseased cell lines, high- and medium-confidence essential genes in healthy *C. elegans* had significantly lower G_C_ than the non-essential genes in the tested pool (Table 1, Table S4).

**Table 1.** *C. elegans* essential genes have significantly lower G_C_ than non-essential genes. (a) Numbers in this column are smaller than in column C because genes with G_C_ equal to zero were excluded from the analysis. (b) Untested includes genes that, possibly due to strong effects on health and/or development, prevented us from obtaining large enough populations of worms for quantitative analyses. This hypothesis, and the observations that led us to propose it, are in agreement with the low G_C_ – essentiality correlation p value observed for this class. (c) Untested corresponds to core metabolic genes that were not tested due to lack of RNAi clone/s or other technical limitations.

| Geneset | Geneset definition | Number of genes in the class | Number of genes with $G_C > 0$ <sup>(a)</sup> | Sign test (median $G_C^{\text{class}} < \text{median } G_C^{\text{all}}$) |
| --- | --- | --- | --- | --- |
| G1 | High-confidence essential | 48 | 38 | $5.808 \times 10^{-5}$ |
| G2 | Medium-confidence essential | 64 | 49 | 0.0427 |
| G1 + G2 | | 112 | 87 | $4.5304 \times 10^{-5}$ |
| G3 | Wild-type | 1095 | 532 | 0.9814 |
| G4 | Untested<sup>(b)</sup> | 174 | 97 | 0.0335 |
| G5 | Unknown<sup>(c)</sup> | 94 | 58 | 0.347 |

In summary, the data show that low G_C_ is associated with essentiality. Given the strong selective pressure on essential genes, and inferring a similar association in organisms beyond these two mammalian species and a nematode, we hypothesize that the property of essentiality may be conserved across taxa for housekeeping genes defined using the G_C_.

### Housekeeping genes preserve organism-specific information

One way to test whether the properties of housekeeping genes defined by having low G_C_ are conserved across evolution is to compare whether the genes with low G_C_ in one species significantly overlap with the low G_C_ genes in another species. Importantly, conservation is the last of the four properties commonly assigned to housekeeping genes; yet not systematically tested.

Thus, we analyzed our low G_C_ genes in the context of a previously published transcriptomics study of nine endothermal organisms^33^ that include chicken, platypus, opossum, mouse, macaque, orangutan, gorilla, chimpanzee, and humans. Since most of these organisms do not have a Gene Ontology available, we analyzed the 1:1 orthologs across all these organisms, which corresponded to 5423 genes. Another advantage of limiting the comparison to the 5423 genes is that it removes the bias in the statistical tests otherwise introduced by the sample size.

Different from the analyses of human datasets, we found that the correlations among the nine endothermic species were lower, and the Spearman correlations of the G_C_ of the 1:1 orthologs distinctively cluster (Figure 5A and 5B). Gini coefficients accurately clustered non-primate mammals distinctively from primates; thus, reinforcing the value of Gini genes not only to identify conserved housekeeping genes but also distinctive biology.

**Figure 5.**
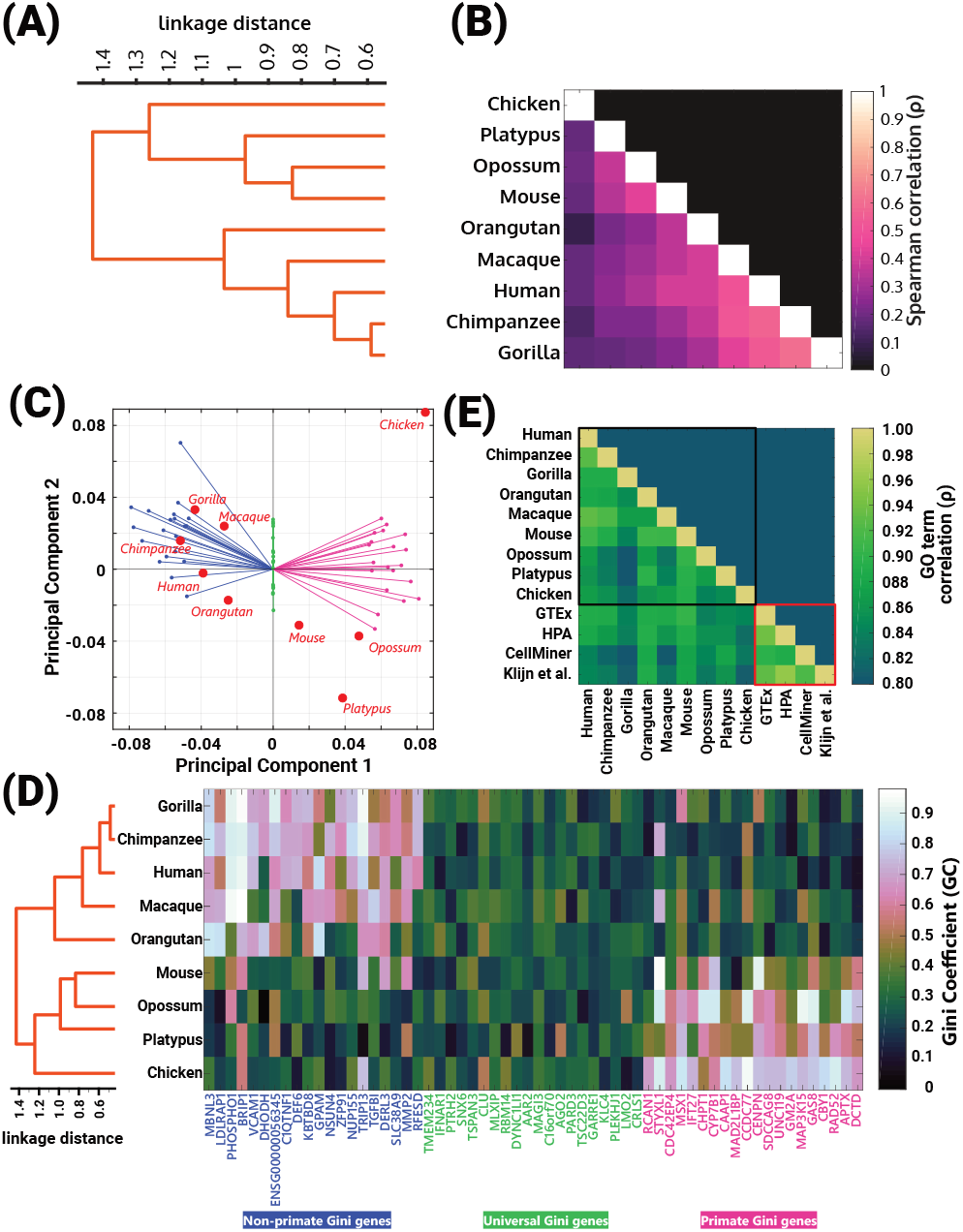
Gini coefficients accurately capture organism-specific differences. (A-B) Jaccard similarity between Gini genes identified using organism-specific transcriptomes capture cluster containing primates. The number of Gini genes with 1:1 orthologs in all organisms is shown using the bar plot on the right of the dendrogram. (C) Principal component 1 (PC1) also captures the cluster containing primates. Also shown are the top 20 primate Gini genes (pink), the top 20 shared Gini genes (green), and top 20 Gini genes in all non-primates (blue) using the principal component coefficients of the first principal components. (D) Correlation among Gini coefficients across different organisms reproduce cluster containing primates (left panel). The Gini coefficients of genes belonging to top 20, middle 20, and bottom 20 coefficients of PC1 are shown (right panel). Left 20 Gini genes are specific to non-primates, middle 20 Gini genes are shared, and right 20 Gini genes are specific to primates. (E) GO term coverage is highly correlated across different datasets, also shown are the GO term correlations with human datasets used in Fig. S6.

Principal Component Analysis (PCA) of G_C_ for 1:1 orthologs was able to reproduce the cluster consisting of primates and the cluster consisting of other organisms (Figure 5C, dendrogram) using the across taxa correlations among Gini genes. The first two principal components accounted for 45.4% of the explained variation (Fig. S5). The first principal component separated the primate cluster from the remaining species. Our clustering also revealed top candidate genes responsible (based on their G_C_ values) for such a clustering. The genes with stable expression across all species (Figure 5D, green), primates only (Figure 5D, pink), or non-primates only (Figure 5D, blue). Interestingly, despite the taxa-specific biology, the coverage of GO terms (Table S2) associated with the Gini genes was highly correlated across all the organisms (Figure 5E (black box)). Together, the results show that individual Gini genes contain important information about species-specific biology, yet higher-level features –such as GO terms– are shared by Gini genes across evolution.

## Discussion

Historically, housekeeping genes have been defined as genes that are consistently expressed across tissues, essential, belonging to cellular maintenance, and conserved across species. Extending from this definition, genes qualified as “housekeeping” genes have been extensively used for benchmarking and normalizing gene expression results in diverse experimental settings, including qRT-PCR, bulk and single-cell transcriptomics, *in situ* hybridization, western blots, FACS, etc. This has led to usage of genes such as Glyceraldehyde 3-phosphate dehydrogenase (GAPDH) becoming commonly used housekeeping genes. Interestingly, our analyses of GAPDH found large differences in its G_C_ values across different datasets (Fig. S7). Thus, we set to use Gini coefficient (G_C_) as a statistical metric to identify inequality in gene expression across different samples.

The Gini coefficient (G_C_) varies between 0 and 1. It quantifies consistency in expression of a gene such that genes expressed in a few conditions or sample types will have a very high G_C_ (closer to 1) while housekeeping genes, by definition, will have lower G_C_ (closer to 0). However, the threshold G_C_ value below which the genes can be classified as housekeeping has not yet been established. The G_C_ value for any given gene depends on its level of expression, but expression depends on the context (cell type, age, stressors etc). Since it is impossible to enumerate all the possible environments an organism, and hence its cells, will encounter, it is impossible to conclusively find gene expression patterns, and unambiguous G_C_ values to define “housekeepingness”. Thus, a challenge going forward lies in identifying G_C_ thresholds or other formulations which can be used to calculate an exact list of housekeeping genes for a given organism. Nevertheless, our comprehensive analyses yielded experimentally supported lists of Gini genes (genes with low G_C_) for different species (Table S3) and, most importantly, we here formalize, for the first time, the definition of housekeeping gene.

In addition to being stable expressed, housekeeping genes have been axiomatically defined as involved in basic cellular functions and essential. Here we use experimental data to test these notions, and show that indeed Gini genes are enriched in GO terms related to cellular maintenance, and are more likely to be essential. Interestingly, for cancer cells, neither the essential genes nor the enriched GO terms fully overlap with those of healthy tissues of the same species suggesting the definition of housekeeping gene validated here can help distinguish health from disease status and possibly aid in the identification of targets for the treatment of disease. Specifically, our comparison of GO enrichment between healthy human tissues and cancer cell lines highlighted the shift from regeneration of cells in healthy tissue to generation of new cells in cancer cells. Further, our analyses found that coverage of GO terms that were distinctly enriched in only tissue datasets was lower than those for cancer datasets (Fig. S4). Thus, rather than cancer cells undergoing dysregulation of certain housekeeping genes, our analysis suggests that cancer has a unique set of housekeeping genes that differs from the one in healthy tissues. Further, these “cancer-enriched” housekeeping genes are associated to cellular division or cellular growth. This contrasts with the notion that housekeeping genes belong to cellular maintenance; and therefore, this criterion may need to be relaxed when defining housekeeping genes. Our analyses also suggest that cellular maintenance (non-growth needs of the cell) and the growth requirements for cellular division are complementary to one another; and their distinction may not lie in the consistency of expression of a gene.

Another axiomatically accepted property of housekeeping genes is that they are “housekeeping” across species. Our analyses using multi-organism datasets showed high GO term correlation across organisms, suggesting conservation of housekeeping pathways, but no significant conservation of individual housekeeping genes. Further, even for housekeeping pathways, the correlation across species is slightly reduced when compared to the correlation across human datasets (Figure 5E). It is possible that this reduced correlation is due to data limitations. The pathway analyses presented here were done using human GO terms; therefore, genes from any given organism were mapped to the corresponding human orthologs and gene identifiers. In fact, the 1:1 ortholog-based GO term analysis used here resulted in eliminating, on average, 69% of the Gini genes from each of the 9 organisms analyzed. Therefore, availability of gene ontology beyond model organisms can provide molecular insight into species-specific biology. Nevertheless, Gc was able to capture taxa-specific biology.

Until now, the concept of housekeeping was often described using a list of genes. As discussed above, in the framework of G_C_, selecting housekeeping genes would require thresholding such that genes which have a G_C_ below a certain value may be regarded as housekeeping genes. However, thresholding eliminates possibly meaningful information. Indeed, our PCA of genes showed that the genes with the highest variation in expression (39.5%) did not enable clustering the 9 organisms (Fig. S8), suggesting that many of the eliminated genes may have critical functions conserved across species. In this study, we also show that even though fewer GO terms were enriched in all the datasets or organisms, the coverage of GO terms that were enriched in each dataset was highly correlated across datasets. These results suggest that housekeeping functions, rather than a list of genes, better described the state of the organism. This explanation has been suggested previously^44^. To test such a hypothesis, there would be a need to generate computer-based models of these organisms that include multiple levels of regulation. Then, one could possibly test *in silico* the expression of genes across different tissues in different contexts (e.g. reduced dietary protein or infection), and infer which genes show the least variation in expression across conditions (low Gc value). Indeed, this means there is a need for functional models for diverse organisms^45,46^.

Key molecular similarities likely underlie the physiological similarities between related species. By crossing Gini coefficients with CRISPR-Cas9 essentiality screens and GO terms we may have captured some of these key molecular similarities as our analysis was able to distinguish primate from non-primate endotherms. On the other hand, even though animals seem phenotypically very different they share molecular similarities that we can capture at the level of GO terms, even if not at the level of specific gene IDs. Nevertheless, what is essential across environmental contexts and taxonomic groups, if anything, is worth future investigation.

Our study only scratches the surface of the answer to practical and fundamental questions and shows the need for organism-specific tools and models; but not just for model organisms, we need models for a diverse set of organisms. Our study suggests that analysis of the ever-increasing “omics” datasets presents an opportunity for better understanding of the biological functions fundamental to sustain life and drive evolution.

## Method

### Literature search

We performed a literature search using Harzing’s Publish or Perish 7^47^ to extract the top 1000 hits from Google Scholar for the query keywords: housekeeping, genes, maintenance, and required. The list of top 1000 papers was downloaded to an excel sheet for further analysis and visualization on MATLAB.

### Data extraction

Transcriptomic datasets were obtained from various sources (Table 2). To resolve differences in gene identifiers, we mapped all to NCBI Entrez gene identifiers using BioMart, within the Ensembl website. When genes did not map to an NCBI gene identifier, we discarded these genes from the analyses.

**Table 2.** Data sources used for this study.

| Organism (sample type) | Data source | Modifications |
| --- | --- | --- |
| **Transcriptomics** |  |  |
| Human (tissues) | HPA<sup>22</sup> | - |
|  | Brawand et al., 2011<sup>33</sup> | Converted to TPM from read per base |
|  | GTEX<sup>34</sup> |  |
| Human (NCI-60 cancer cell lines) | CellMiner<sup>48</sup> | - |
|  | Klijn et al. 2015<sup>35</sup> | - |
| *C. elegans* (cell types) | Cao et al., 2017<sup>18</sup> | - |
| Chicken, Platypus, Orangutan, Bonobo, Gorilla, Chimpanzee, Macaque, Mouse, Opossum (tissues) | Brawand et al., 2011<sup>33</sup> | Converted to TPM from read per base |
| Chinese hamster (tissues) | Shamie I.S.*, Duttke S.H.*, la Cour Karottki K.J., Han C.Z., Hansen A.H., Hefzi H., Xiong K., Li S., Roth S., Tao J., Lee G.M., Glass C.K., Kildegaard H.F., Benner C., Lewis N.E. A Chinese hamster transcription start site atlas that enables targeted editing of CHO cells. bioRxiv, (2020). DOI: 10.1101/2020.10.09.334045<sup>49</sup> |  |
| Chinese hamster ovaries (cell lines) | See Table S6 for accession IDs |  |
| **Essentiality screens** |  |  |
| Human (NCI-60 cancer cell lines) – CRISPR-Cas9 | DepMap<sup>38–40</sup> | - |
| CHO cell lines – CRISPR-Cas9 | Xiong et al. 2020<sup>37</sup> and Table S5 | - |
| *C. elegans* (cell types) - RNAi | Unpublished study provided by Eyleen J. O'Rourke; method described in Ke et al. 2018<sup>50</sup> and Supplementary text. | - |

### Gini Coefficient (G_C_)

The G_C_ measures the inequality in frequency distribution of a given parameter (e.g., levels of income, income mobility^51^, education^52^, etc.) compared to the frequency distribution of total population^32^. For analysis of transcriptomic data, the parameter is expression of a given gene and is compared against the total gene expression is distributed across different samples^30^. The G_C_ is calculated as the ratio of area between the Lorenz curve and line of equality over the total area under the line of equality. The Lorenz curve is the graphical representation of the distribution of a given parameter; and is given by eqn. (1):

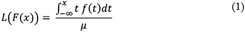

where *μ* denotes the average, *f*(*x*) denotes the probability density function, and *F*(*x*) denotes the cumulative distribution function. The calculation was implemented in MATLAB (R2016b), for which the code is available at GitHub (https://github.com/LewisLabUCSD/gene-gini-matlab).

### Gene Ontology (GO) enrichment

Due to lack of availability of unique gene ontologies for the different organisms discussed in the study, genes of the organisms that mapped to the human ortholog genes were used to identify the respective GO term. Here, hypergeometric tests were used to check whether the number of genes associated to a GO term, in the query list, are more significant given the distribution among GO terms in the subject gene list. GO terms associated to human genes were downloaded from Gene Ontology Consortium webpage (http://current.geneontology.org/products/pages/downloads.html). All analysis was focused only on the Biological Process (P) aspect. All p-values were calculated using hypergeometric test for overrepresentation reported after correction using the Benjamini Hochberg FDR. A GO-term was considered enriched if the corrected p-values were below 0.05.

