## Supplementary for "What are housekeeping genes?"

### Supplementary Text

#### Supplementary results

##### Definition of housekeeping genes lacks experimental basis

We scoured the literature from Google Scholar using Herzing's Publish or Perish[1] software. From the top hits with highest citations among more than 109,000 search results, we looked at the text and title of articles containing the words: housekeeping, genes, maintenance, and required. Excerpts from the most highly cited articles have been listed in the Supplementary text. We found that none of these articles reference an article which constructs the experimental basis of the definition of housekeeping genes. This suggests that, given the pervasiveness and usefulness of the term, there is a need to provide an experimental basis for existence of housekeeping genes. The list of definitions used in some of the highly referenced papers is below:

1. Thellin, Zorzi, Lakaye, De Borman, Coumans, Hennen, Grisar, Igout, and Heinen, 1999 (Cited 1761) – "All these techniques can use internal standards, mainly housekeeping genes, so called because their synthesis occurs in all nucleated cell types since they are necessary for the cell survival."[2]

2. Eisenberg and Levanon, Trends in Genetics, 2003 (Cited – 754) - "… most genes show constitutive expression in only a subset of tissues, some gene products are required for the maintenance of the basal cellular function and are constitutively found in all human cells. These genes are called housekeeping genes."[3]

3. Eisenberg and Levanon, Trends in Genetics, 2013 (Cited – 726) – "Housekeeping genes are involved in basic cell maintenance and, therefore, are expected to maintain constant expression levels in all cells and conditions." "Housekeeping genes are genes that are required for the maintenance of basal cellular functions that are essential for the existence of a cell, regardless of its specific role in the tissue or organism."[4]

4. Warrington, Nair, Mahadevappa, and Tsyganskaya, Physiological Genomics, 2000 (Cited - 603) – "housekeeping genes, or maintenance genes, are those genes constitutively expressed to maintain cellular function."[5]

5. Butte, Dzau, and Glueck, Physiological Genomics, 2001 (Cited – 212) – "Housekeeping genes are constitutively expressed to maintain cellular function."[6]

6. Zhu, He, Hu, and Yu, Trends in Genetics, 2001 (Cited - 260) – "Housekeeping (HK) genes are ubiquitously expressed in all tissue and cell types and constitute the basal transcriptome for the maintenance of basic cellular functions."[7]

While we did not find an experimental basis from literature search, we did find a commonality. The housekeeping genes appeared to delineate relationship between three types of biological features: (i) consistently or stably expressed genes, (ii) essential genes, and (iii) genes participating in cellular maintenance. In the subsequent sections, we will test multiple different datasets belonging to different organisms to establish relationship between these biological features.

##### GAPDH may not be a good candidate as a housekeeping gene

Glyceraldehyde 3-phosphate dehydrogenase (GAPDH) is the most commonly used housekeeping gene to benchmark expression of other genes in qRT-PCR analyses. To define the appropriateness of GAPDH as a housekeeping gene, we started with previously published transcriptomics data belonging to CHO cells, hamster tissues[8], human tissues from Genotype-Tissue Expression (GTEx) project[9] and Human Protein Atlas (HPA)[10], and NCI-60 cancer cells (Klijn et al.[11], and CellMiner[12]). We calculated the $G_C$ for these datasets and then compared the $G_C$ of GAPDH across the 7 datasets.

Low $G_C$ represents low variability in level of expression across tissues or samples, as would be expected for housekeeping genes. However, our analyses indicated that the $G_C$ values for GAPDH were very different across all human and hamster datasets. For instance, Klijn et al. (i.e., NCI-60 cell lines) showed the lowest $G_C$ value was at 14.6 percentile and hamster tissue had the highest $G_C$ value at 57.6 percentile (Figure S1). The $G_C$ values in human datasets varied by 18 percentiles while hamster and CHO data differed by 35.3 percentiles. Thus, the high variability in $G_C$ percentiles indicate that GAPDH may not be a good candidate as a housekeeping gene as it is not as stably expressed as generally thought.

### Supplementary Methods

#### Whole animal *C. elegans* RNAi screens

*C. elegans* rrf-3 mutants, which are hypersensitive to RNAi, were used. The details on how the RNAi worms were grown and prepared can be found in Ke et al. 2018[13]. Please see below for details on manual screening.

##### *Workflow for scoring*

1. Open 1-2 L4440 representative reference images, leave them open in 2nd monitor.
2. Open well image to be scored.
3. Compare to reference images and decide whether size of worms is = or < than worms in reference image. Enter score in column “reduced size” (0 = WT, 1= reduced size).
4. Define whether smaller animals are gravids (small but have eggs) or larvae (small but no eggs): (i) Enter score in “gravids” column (1=gravids, 0=larvae), OR (ii) Enter score in "Larvae or egg less adult" column (0=gravids, 1=Larvae or egg less adult).
5. A second round of sorting was done based on the criterion described as follows: The sum of the scores from the column "reduced size" (which can be gravid with reduced size or larvae) was calculated. The maximum score could be 6 (3 repeats of the experiment, each scored independently by 2 personnel) or 4 if there were only 2 repeats of the experiment.
6. The genes were grouped based on the following criteria:
   <u>Group 1: High confidence hit</u>
   RNAi clone consistently leading to reduced size (it can be larvae or adult with reduced size).
   For plates tested in triplicate: High confidence hits are those RNAi clones that scored 5 or 6 (out of 6 scores). This means that in all 3 repeats worms had reduced size (result was very reproducible).
   For plates tested in duplicate: High confidence hits are those RNAi clones that scored 4. This means that the worms had reduced size in both the repeats.
   <u>Group 2: Medium confidence hit</u>
   For plates tested in triplicate: Medium confidence hits are those RNAi clones that scored 4. This means that in 2 repeats out of 3, worms had reduced size.
   For plates tested in duplicate: Medium confidence hits are those RNAi clones that scored 3 (out of 4 scores).
   <u>Group 3: Wild Type BIOMASS</u>
   RNAi clones that scored 0-1 (in plates that were done in duplicate or triplicate).
   <u>Group 4: Untested</u>
   RNAi clones for which bacteria do not grow. Hence, worms in these wells are starved.

RNAi clones that did not grow in 2 out of 3 or 1 out of 2 trials,
Due to experimental issues, these clones cannot be called neither Wild Type nor reduced size. If these clones were critical to the model, they could be manually retested.
Group 5 Unknown
RNAi clones that have given variable results (scores 2-3), therefore we cannot say with confidence whether they are WT or show reduced size.

#### Supplementary Table Captions

**Table S1. GO term enrichment analysis of Gini genes identified using human tissues and cancer cell line datasets.** Shown are 1244 terms enriched in at least one of the following human datasets – GTEx, HPA, CellMiner, Klijn et al.

**Table S2. GO term enrichment analysis of Gini genes identified using transcriptomes published in Brawand et al.** Shown are 918 terms enriched in at least one of the organisms. The results only account for genes which have a 1:1 orthologs in all the organisms.

**Table S3. List of Gini genes identified using each of the 15 transcriptomes -** GTEx, HPA, CellMiner, Klijn et al., 9 organisms in Brawand et al., *C. elegans* cell types, hamster, and CHO cells. Each tab of the excel file is one transcriptome.

**Table S4. List of C. elegans genes in each of the 5 groups classified using RNAi screens.**

**Table S5. List of essential genes in CHO cells as identified by Kai et al. 2020**[14]

**Table S6. List of accession IDs from where CHO transcriptomics were assembled**

#### Supplementary Figures

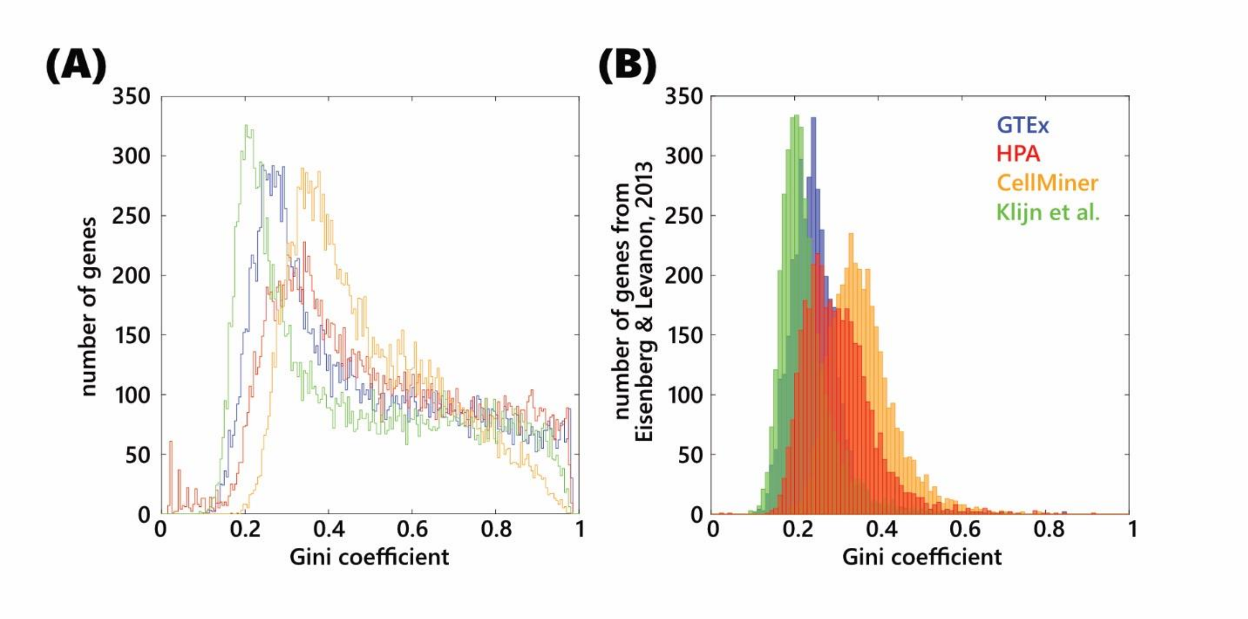


**Figure S1. List of housekeeping genes published by Eisenberg & Levanon have low Gini coefficients when using GTEx, HPA, CellMiner, or Klijn et al. datasets.** (A) The distribution of the Gini coefficients of all the genes for each of the datasets. (B) The distribution of the Gini coefficients of only the list of genes published by Eisenberg & Levanon.

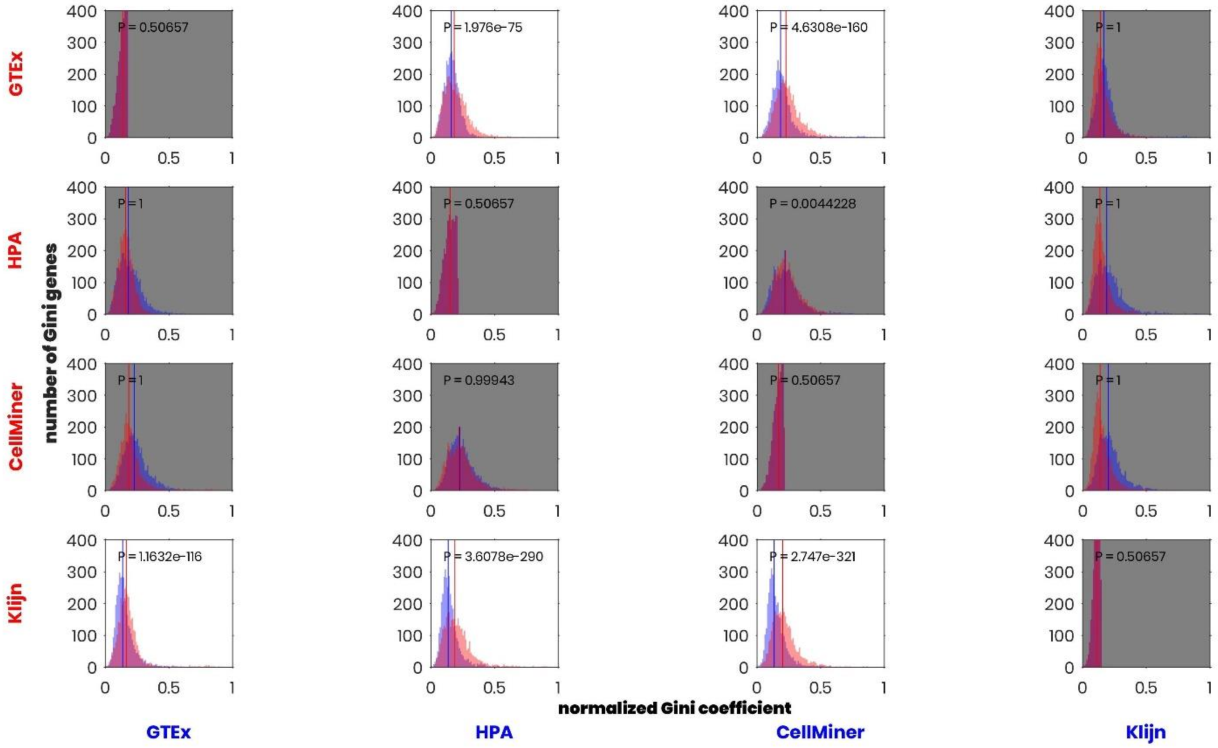


**Figure S2. Cross comparison of Gini genes across datasets.** Gini genes identified using one dataset had lower Gini coefficient in the other dataset. The grey background plots are those where the P-value calculated suing Wilcoxon rank sum test was not significant (p <0.00001).

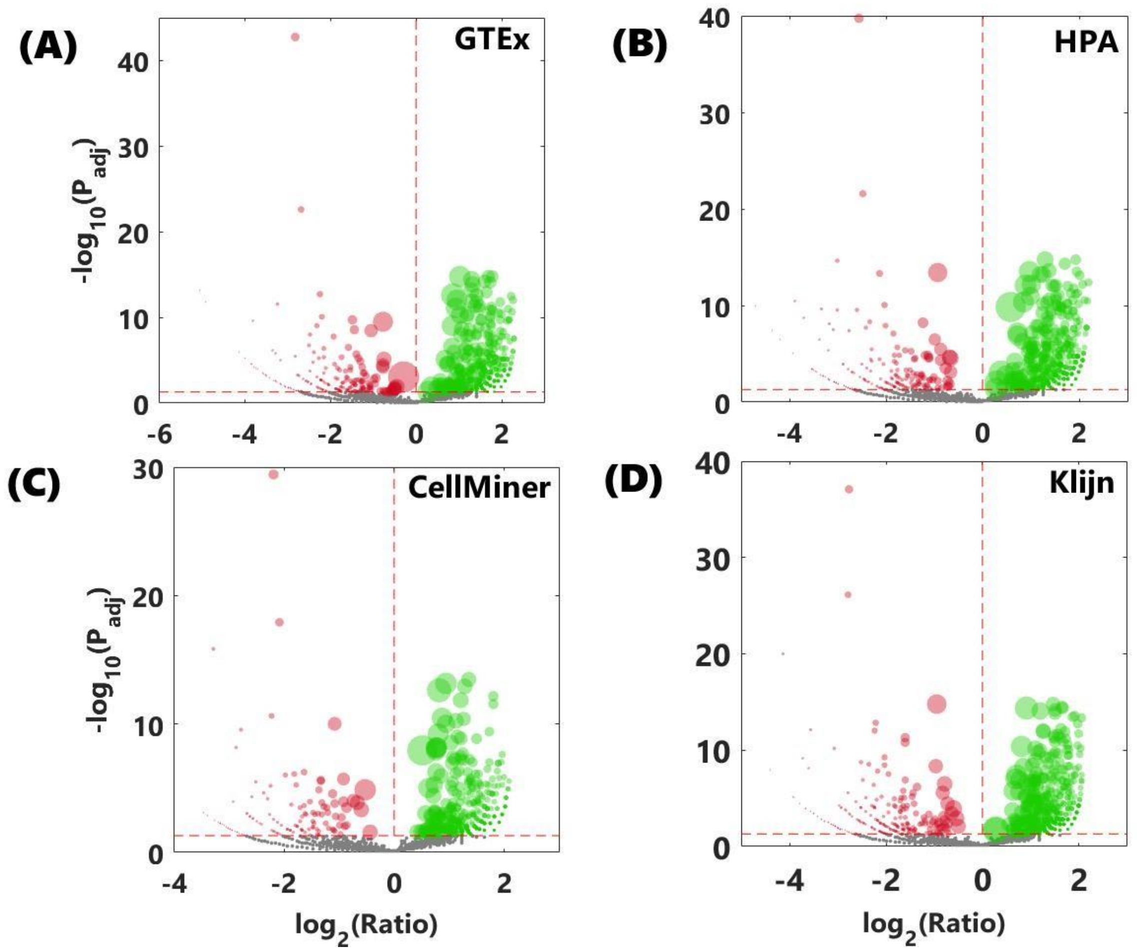


**Figure S3. Volcano plots for GO term enrichment analysis for GTEx (A), HPA (B), CellMiner (C), and Klijn et al. (D).** The colors green, red, and grey indicate the GO terms which were over-represented, under-represented, and not enriched. The size of the bubble indicates number of genes belonging to a GO term. The x-axis represents the ratio of number of hits in the list of Gini genes to those achieved at random.

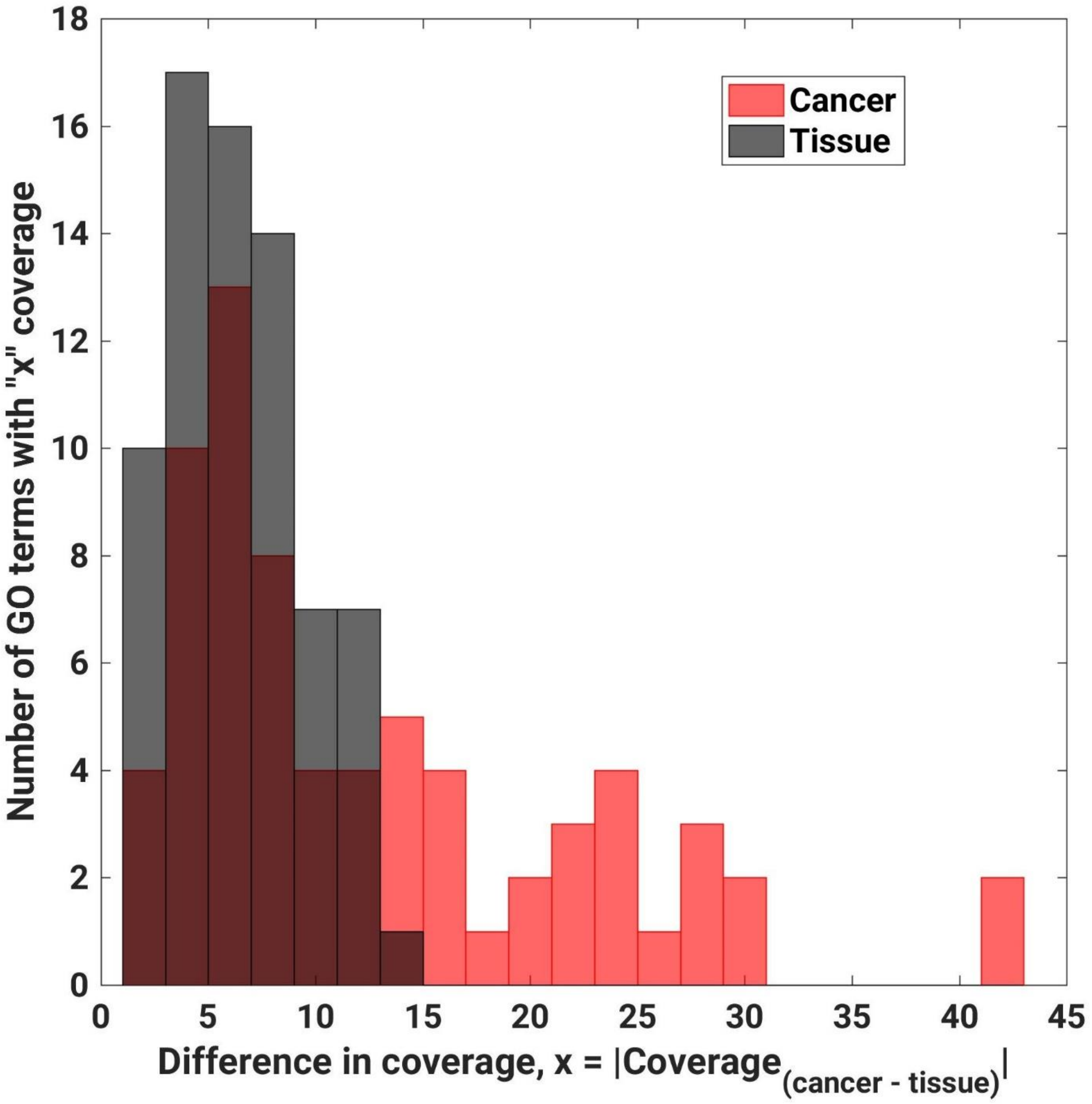


**Figure S4. Coverage of GO terms enriched in only cancer datasets was higher than coverage of GO terms enriched in only tissue datasets.** Red bars for cancer cells and Black bars are for tissues. The full data used to prepare this plot is in Table S1. Included in this plot are 70 GO terms which are only enriched in cancer datasets but in neither of the tissue datasets, and 77 GO terms which are only enriched in tissue datasets but in neither of the cancer datasets.

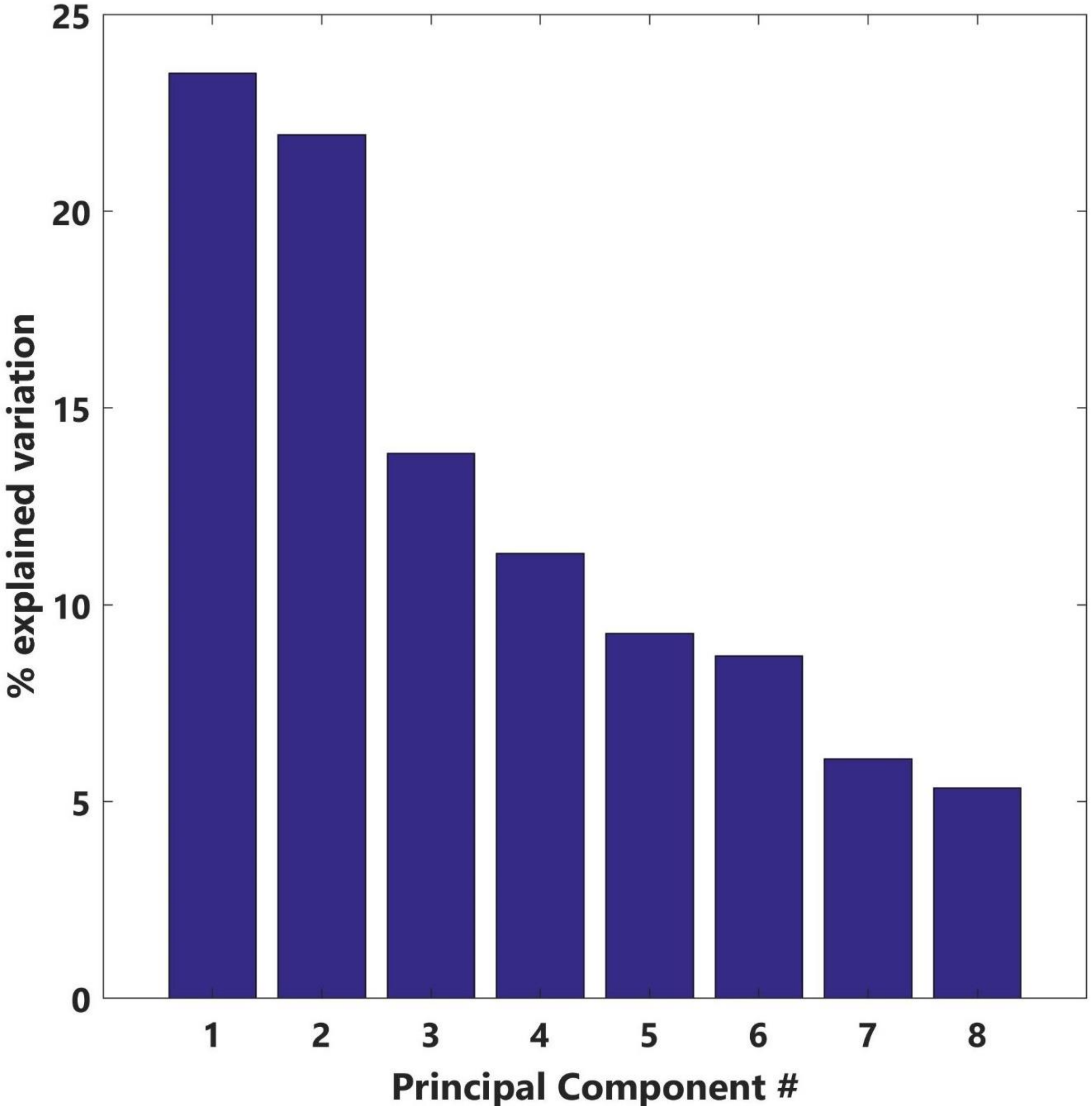


**Figure S5. Percentage of explained variation in Principal Component Analysis.** The PCA was performed using 1:1 ortholog Gini coefficients which were calculated using transcriptomes in Brawand et al. to cluster organisms.

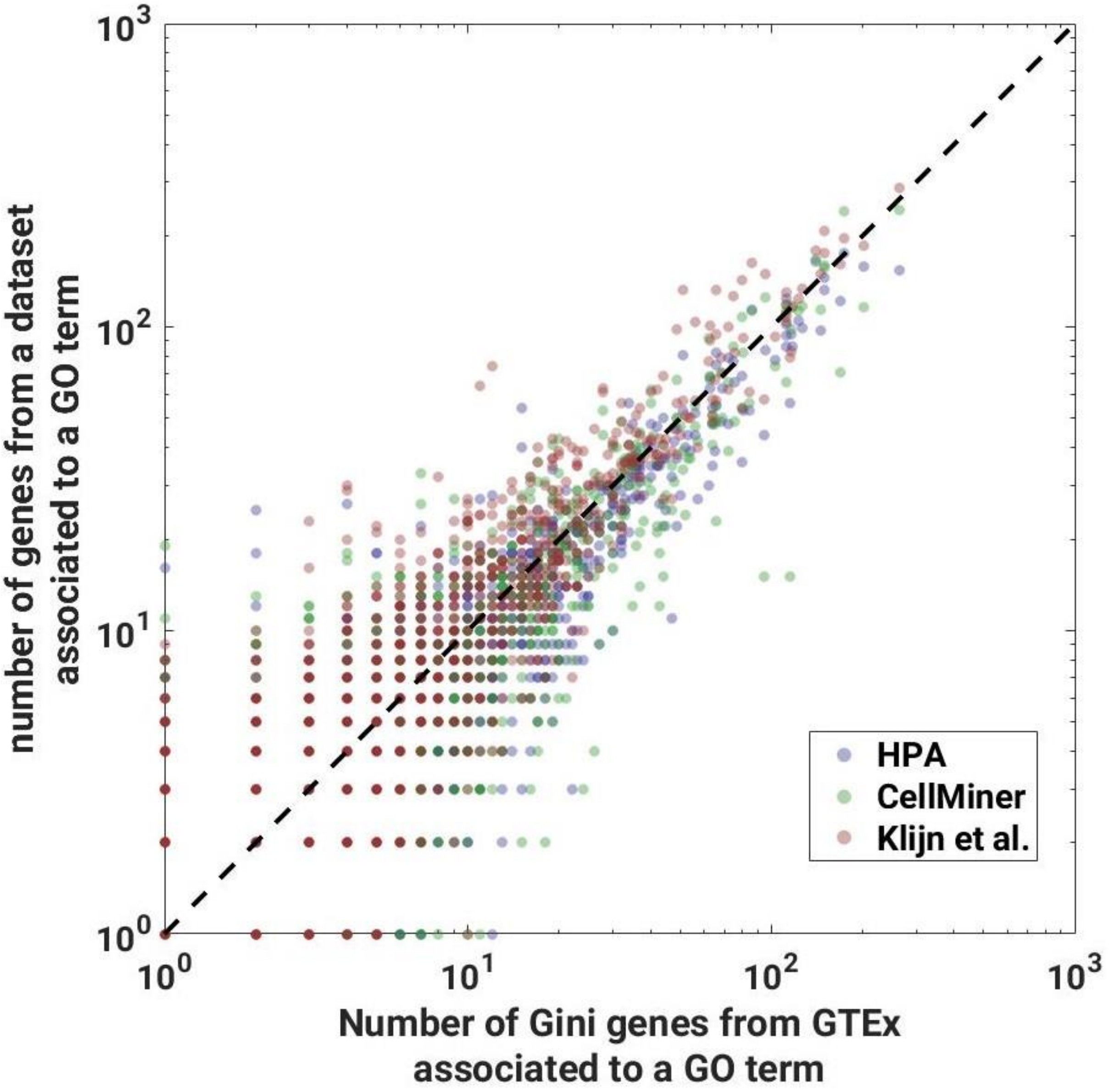


**Figure S6. GO term coverage is highly correlated across human datasets.** HPA/GTEx comparison represented in blue, CellMiner/GTEx in green, and Klijn/GTEx in red.

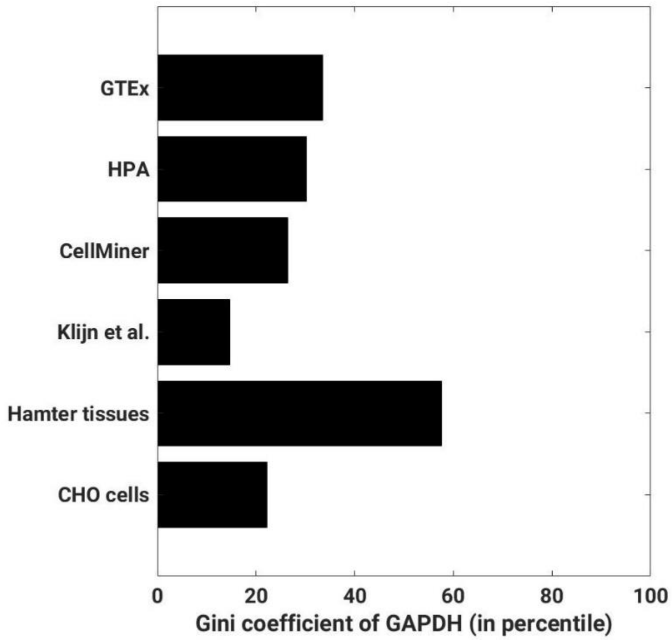


**Figure S7. Glyceraldhyde 3-phosphate dehydrogenase (GAPDH) may not be a good choice for housekeeping gene.** Gini coefficients were converted to percentiles (x-axis) using each of the datasets (y-axis). GAPDH has high Gini coefficient in most of the datasets.

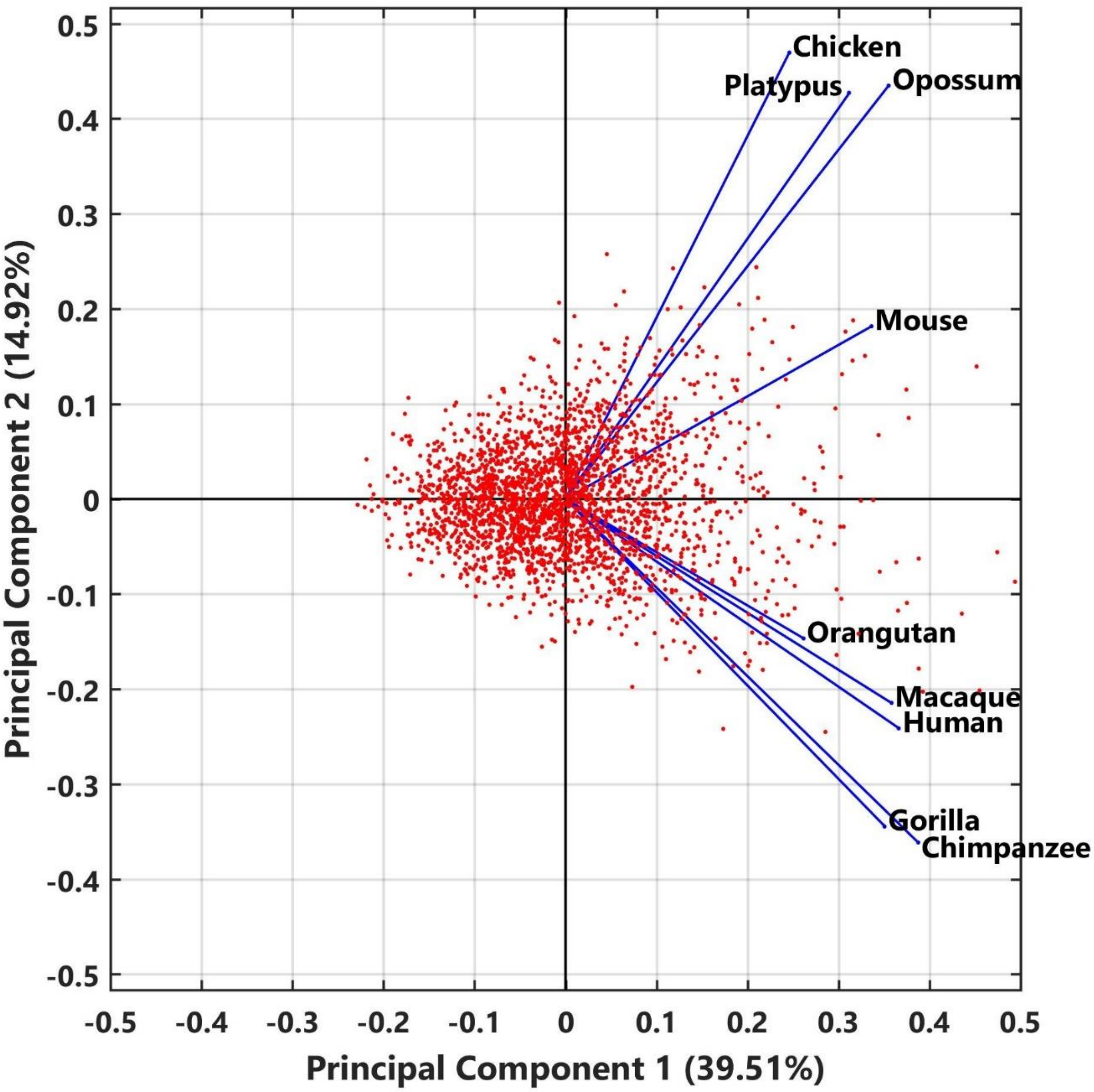


**Figure S8. A scatter plot showing principal components 1 & 2 if genes and organisms switched places in the analysis.** The first principal component which captures majority of explained variation does not explain Gini values in either of the organisms as all of them lie to the right-hand side of the plot.
